# Cerebellar volume in autism: Meta-analysis and analysis of the ABIDE cohort

**DOI:** 10.1101/104984

**Authors:** Nicolas Traut, Anita Beggiato, Thomas Bourgeron, Richard Delorme, Laure Rondi-Reig, Anne-Lise Paradis, Roberto Toro

## Abstract

Cerebellar volume abnormalities have been often suggested as a possible endophenotype for autism spectrum disorder (ASD). We aimed at objectifying this possible alteration by performing a systematic meta-analysis of the literature, and an analysis of the Autism Brain Imaging Data Exchange (ABIDE) cohort. Our meta-analysis sought to determine a combined effect size of ASD diagnosis on different measures of the cerebellar anatomy, as well as the effect of possible factors of variability across studies. We then analysed the cerebellar volume of 328 patients and 353 controls from the ABIDE project. The meta-analysis of the literature suggested a weak but significant association between ASD diagnosis and increased cerebellar volume (p=0.049, uncorrected), but the analysis of ABIDE did not show any relationship. The studies in the literature were generally underpowered, however, the number of statistically significant findings was larger than expected. Although we could not provide a conclusive explanation for this excess of significant findings, our analyses would suggest publication bias as a possible reason. Finally, age, sex and IQ were important sources of cerebellar volume variability, however, independent of autism diagnosis.

## Introduction

Autism spectrum disorders (ASD) affect 1% of the population and are characterised by impairments in social interactions, and the variety of interests. Through the years, many reports have suggested that cerebellar abnormalities may be implicated in the onset of Autism spectrum disorders (ASD) (review in Wang, Kloth, and Badura 2014). The cerebellum is a heavily folded region of the rhombencephalon, with a number of neurons comparable to those of the neocortex. It is characterised by a highly regular arrangement of neurons and connections, supposed to support massive parallel computing capabilities in particular through long term synaptic plasticity (D’Angelo 2014; Dean et al. 2010)). The cerebellum has been traditionally involved in the performance of precise motor behaviour, and patients with ASD also present varying degrees of dyspraxia (Dziuk et al. 2007; Åhsgren et al. 2005). There is also growing evidence for an involvement of the cerebellum in cognitive and affective functions, which could be impaired in autism (Leiner, Leiner, and Dow 1993; Manni and Petrosini 2004; Rondi-Reig et al. 2014; Schmahmann and Sherman 1998).

The first case report of abnormal cerebellar anatomy in autism was published in 1980 (Williams et al. 1980), and described a reduced count of Purkinje cells in the cerebellar vermis of a patient with ASD. In 1987, Courchesne et al. were the first to find a cerebellar abnormality in a patient with ASD using in vivo magnetic resonance imaging (Eric Courchesne et al. 1987). Since then, various studies comparing the volumes and areas of cerebellum sub-regions between patients with ASD and controls with typical development have described significant differences. These studies pointed at the cerebellar volume as an interesting endophenotype for ASD: the volume of the cerebellum and its different subregions being relatively easy to measure from standard T1-weighted magnetic resonance imaging data. However, whereas many articles report statistically significant differences, many others fail to detect such differences. These discrepancies could be due to many factors, for example, to the high heterogeneity in the etiology of ASD, to differences in the inclusion criteria across studies, or to differences in the eventual comorbidities affecting patients in different groups. The discrepancies could also reveal methodological bias, such as differences in MRI sequences, segmentation protocols, or statistical analyses. Finally, they could also result from chance when small sample sizes lead to noisy estimations of mean volumes.

Here, our aim was thus to objectify the alterations of cerebellar volumes in ASD. In the first part of this article we describe our systematic meta-analysis of the literature to examine the differences across previous reports and to determine a combined effect size. In the second part, we describe our analysis of cerebellar volume in the ABIDE cohort (Di Martino et al. 2014), and study the consistency of these results with those from the meta-analysis. Finally, we describe our analyses of the impact of distinct sources of variability such as sex, age at inclusion or IQ on the volume differences between patients with ASD and controls.

## Materials and methods

### 1. Meta-analysis of the literature

#### Collection and selection of articles

We queried PubMed on October 12, 2016 for all the articles that met the search criteria “cerebell* AND autis*”. When one of these results was a systematic review or a meta-analysis on the differences of volumes and surfaces between individuals with ASD and controls on several brain regions including the cerebellum, we added all their cited references.

The selection of articles was made in two steps. First, we screened all the titles and abstracts to eliminate the articles that did not meet the following inclusion criteria: we kept articles written in English with the title or abstract indicating that the size of a brain region was measured in individuals with ASD and we removed items like books, invalid references, reviews, meta-analyses or other studies where the cerebellum was excluded from analysis, as well as studies focused in functional activity, molecules, or with measurements made with technologies other than magnetic resonance imaging.

Second, we recovered the selected articles in their full versions, and included those with available volumetric or area MRI measurements (mean and standard deviation) on at least one region of the cerebellum for both individuals with ASD and healthy controls. Each selected region had to be measured in at least five articles to be included. We tried to avoid the overlap of subjects between studies, favouring articles with more complete information and larger cohorts. We kept the articles regardless of whether individuals with autism exhibited neurological disorders such as epilepsy or followed a pharmacological treatment.

#### Data abstraction

For each selected article we collected means and standard deviations of cerebellar region volumes or areas in order to calculate Cohen's d effect size. When there were more than one ASD or healthy control group, we reconstructed the mean (*x*) and standard deviations (*s*) of the merged group from the means (*x*_*i*_), standard deviations (*s*_*i*_) and numbers of subjects (*n*_*i*_) of the subgroups:

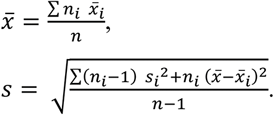

We also collected mean age and Intelligence Quotient (IQ) for analysing their impact on volume or area differences using meta-regression. When such means were not reported, we approximated them from mid-ranges or textual descriptions.

When the volume data were provided for two separate regions, we estimated the mean and the standard deviation of the combined region using the following formulae:

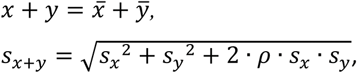
where *ρ* is the assumed correlation coefficient between the two regions. If the two regions in question were two areas of different types, we assumed *ρ* = 0.5.

#### Meta-analysis

We conducted two meta-analyses. The first one compared mean cerebellar volume between ASD patients and controls. Effect sizes were computed a standardised mean differences (Cohen’s d) using Hedges’ g as estimator (Hedges 1981). The second meta-analysis compared the variability of cerebellar volumes between patients and controls by computing the log-variance ratio (Traut 2017). We combined effect sizes using a random effects model. In such a model, if the variability between the estimated effects is larger than expected from sample variance alone, studies are considered heterogeneous, i.e., there is also variability between the true effects (Borenstein et al. 2009, pt. 3). The between-study variance *τ*^*2*^ was estimated using the restricted maximum likelihood (REML) method from which we computed the proportion of variance imputable to heterogeneity (*I*^*2*^). To assess the impact of age and IQ on the effect size and thus identify possible sources of heterogeneity, we also conducted a meta-regression with average age and average IQ of patients with ASD as fixed effects (Borenstein et al. 2009, chap. 20). After the meta-regression, we could estimate an effect size for each study, removing the linear contribution of age and IQ.

We evaluated publication bias and p-hacking in several ways. First, we calculated the rate of studies showing a statistically significant difference between ASD and controls. We compared this rate to the average statistical power obtained by assuming that the actual effect is equal to the effect estimated after the meta-regression for each study. Second, we evaluated the symmetry of the funnel plot (Light and Pillemer 1984) using Egger’s test. Egger’s test relies on the assumption that studies with larger samples are less subject to publication bias than studies with smaller samples and that significant results would tend to report larger effect sizes (Egger et al. 1997). Finally, we plotted the p-curve which shows the distribution of significant p-values. We evaluated p-curves for the inferred power – the most likely statistical power of the studies to get the observed p-curve (Simonsohn, Nelson, and Simmons 2014).

The computations were performed using R (https://www.rproject.org) and the packages meta and metafor along with the online p-curve app 4.0 (http://www.p-curve.com/app4/). We report statistical significance for an alpha level of 0.05. P-values are not corrected for multiple comparisons.

### 2. Analysis of ABIDE

We analysed the MRI data made available by the ABIDE (Autism Brain Imaging Data Exchange) project, which includes 539 brain MRI of individuals with ASD and 573 brain MRI of controls. Cerebellum volumes were automatically segmented using FreeSurfer 5.1. We developed a tool for FreeSurfer segmentations to visually check the quality of the segmentation for each subject (**Figure 1**, https://github.com/r03ert0/QCApp-Cb). We kept only the subjects for whom the segmentation quality was clearly good. Some MRIs with minor segmentation errors could have been manually corrected, however, because of their number we chose to exclude them from the present analyses. Our quality control focused on the cerebellum, but we combined its results with our previous quality control of the whole brain (Lefebvre et al. 2015) and conserved only the subjects that passed both.

**Figure 1.**
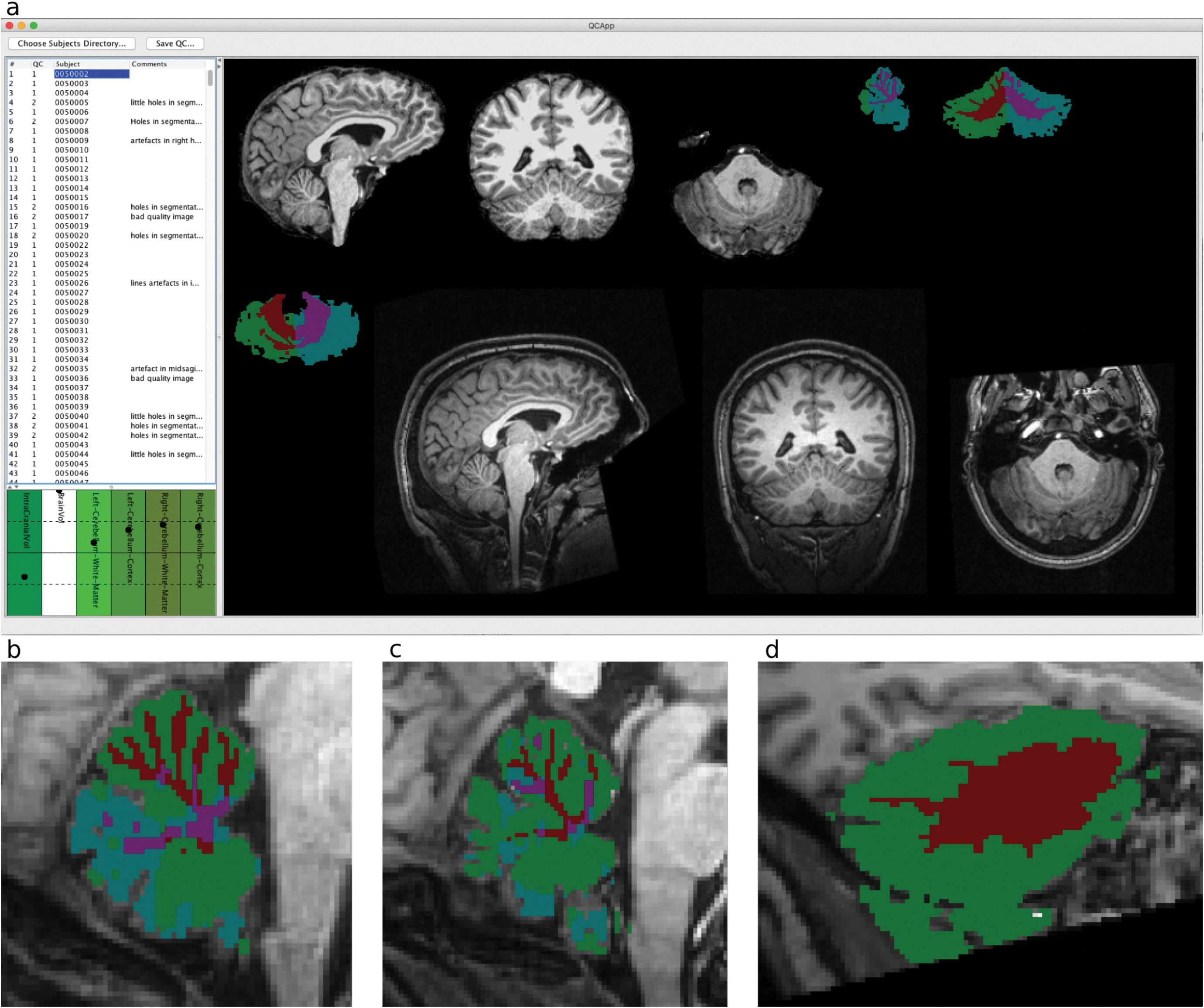
Quality control of the cerebellum automatic segmentation. **a.** Quality control tool: for each subject, we evaluated the quality of the segmentation in comparison with the original image in the three different planes. The values of the different measured volumes along with their deviation from the overall mean in represented in the graph at the bottom left. **b.** A segmentation evaluated as correct; **c.** A segmentation with an excess of unlabelled regions. **d.** A segmentation with major labelling errors.

Following our previous results showing the nonlinear variation of brain anatomy relative to brain volume (Toro et al. 2008; Toro et al. 2009; Lefebvre et al. 2015), we studied the allometric scaling of cerebellar volume. The division of regional volume measurements by total brain volume (“normalisation”) is often used to control for differences in brain volume between groups. This strategy would only be appropriate if the volume of the cerebellum scaled proportionally to total brain volume. We assessed the scaling factor of the cerebellum with total brain volume by regressing the logarithms of total brain volume (BV) and the volume of the cerebellum (Cb):

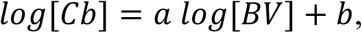
where *a* is the scaling factor.

We evaluated the effect of diagnosis and other factors on whole cerebellum volume, cerebellar white matter volume and cerebellar grey matter volume with two linear models. The first model included group, age, IQ, scanning site, sex and brain volume as fixed effects; the second model included in addition the interactions of group with age, IQ, scanning site, sex and brain volume. Because these statistical models are not the same as those used in the meta-analysis of the literature, we also analysed the ABIDE data using the same meta-analytical approach as for the literature. In the meta-analysis of ABIDE the volume estimations of each site were combined using a random effects model, where each estimation is weighted by the inverse of the estimation’s variance. At each site we eliminated the minimal number of subjects that would ensure that the age and sex matching were respected, as in the meta-analysis of the literature. We did this iteratively, eliminating one subject after another, until the p-values on the differences of age mean, age variance and sex ratio were each above 0.2. We used JMP Pro 12.1.0 for fitting linear models and R version 3.3.1 for the meta-analysis approach.

## Results

### 1. Meta-analysis of the literature

#### Selection of articles

The PubMed queries (“cerebell* AND autis*”) returned 947 items that were combined with the 124 references cited by a systematic review (Brambilla et al. 2003) and a meta-analysis on cerebellum in ASD (Stanfield et al. 2008). We also added two studies (Langen et al. 2009; Evans et al. 2014) that we found by other means. After removing duplicates, we obtained a total of 1029 articles. We screened the titles and abstracts of these items with our relevance criteria and recovered the full text of 76 articles. Our final analyses were based on 30 articles covering seven different regions of interest (see **Figure 2** for the PRISMA workflow). The three most reported measures were whole cerebellum volume (1050 subjects in total), vermal lobules VI-VII areas (965 subjects in total) and vermal lobules I-V areas (861 subjects in total) (see **Table 1** for a description of the selected articles).

**Figure 2.**
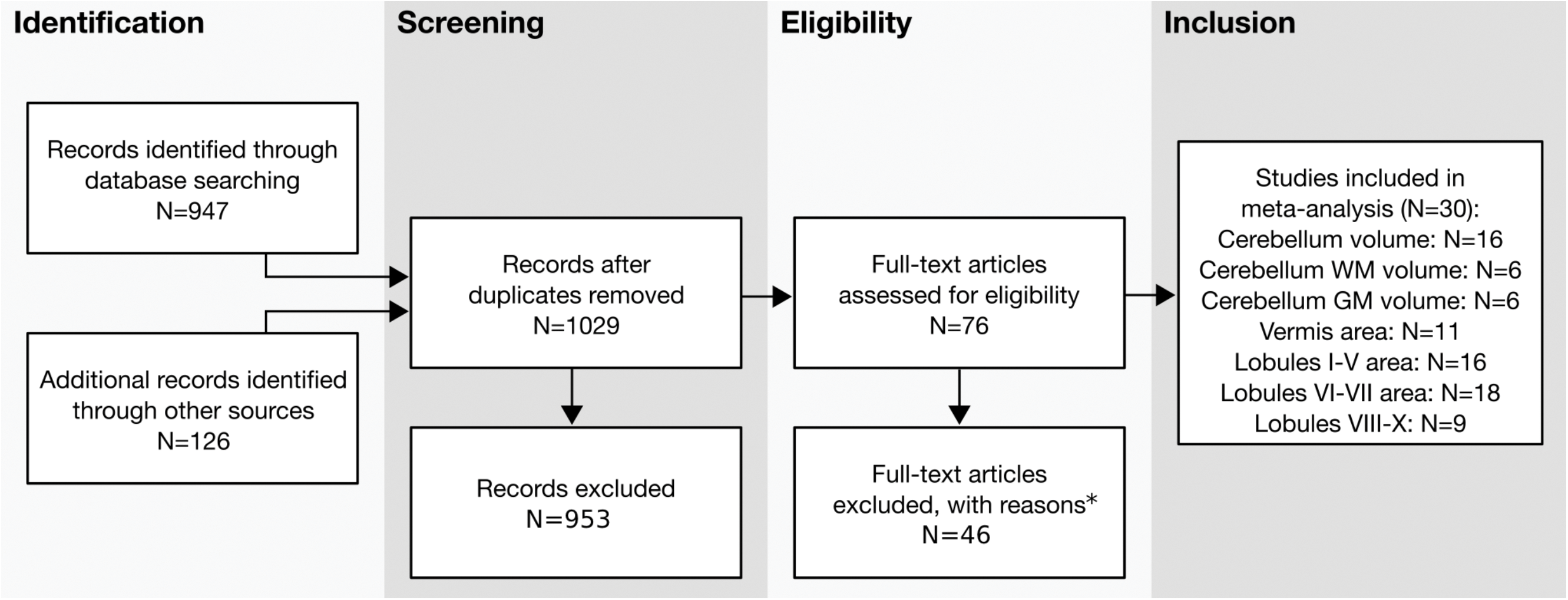
PRISMA workflow for identification and selection of studies (Moher et al. 2009). * no measure on cerebellum: n=7, mean or SD not retrievable: n=16, no typical control group: n=7, no autism group n=1, review or meta-analysis: n=5, subjects included in another study: n=10

**Table 1.** Studies included in the meta-analysis

| Study | N<sub>ASD</sub> (F) | N<sub>Ctrl</sub> (F) | Age<sub>ASD</sub> | Age<sub>Ctrl</sub> | IQ<sub>ASD</sub> | IQ<sub>Ctrl</sub> | Regions of interest |
| --- | --- | --- | --- | --- | --- | --- | --- |
| Hodge et al. 2010 | 22 (0) | 11 (0) | 9.4±2.0 | 10.4±2.7 | 87±22 | 114±11 | Cb, WM, GM, Vermis, I-V, VI-VII, VIII-X |
| Scott et al. 2009 | 48 (0) | 14 (0) | 12.4±3.1 | 12.5±3.1 | 79±23 | 113±12 | Cb, Vermis, I-V, VI-VII, VIII-X |
| Hallahan et al. 2009 | 114 (18) | 60 (7) | 31.9±10.1 | 32.0±9.0 | 97±18 | 114±12 | Cb |
| Webb et al. 2009 | 45 (7) | 26 (8) | 3.9±0.3 | 3.9±0.5 | 59±21 | 115±15<sup>b</sup> | Cb, Vermis, I-V, VI-VII, VIII-X |
| Langen et al. 2009 | 99 (8) | 89 (7) | 12.9±4.5 | 12.4±4.8 | 108±14 | 110±13 | Cb |
| Lahuis et al. 2008 | 21 (0) | 21 (0) | 11.1±2.2 | 10.4±1.8 | 107±14 | 103±15 | Cb |
| Cleavinger et al. 2008 | 28 (0) | 16 (0) | 13.9±5.3 | 13.9±5.4 | 99±18 | 102±14 | Cb, WM, GM, Vermis, I-V, VI-VII, VIII-X |
| Catani et al. 2008 | 15 (0) | 16 (0) | 31.0±9.0 | 35.0±11.0 | 109±17 | 120±21 | Cb |
| Bloss and Courchesne 2007 | 9 (9) | 14 (14) | 3.7±0.9 | 3.8±1.1 | 83±18 | 119±13 | Cb, WM, GM |
| Hazlett et al. 2005 | 51 (5) | 14 (4) | 2.7±0.3 | 2.4±0.4 | 54±9 | 108±19 | Cb, WM, GM |
| Palmen et al. 2004 | 21 (2) | 21 (1) | 20.1±3.1 | 20.3±2.2 | 115±19 | 113±10 | Cb |
| Kates et al. 2004 | 9 (1) | 16 (2) | 7.6±2.4 | 8.3±2.4 | 70±19 | 124±10 | Cb, WM, GM |
| Akshoomoff et al. 2004 | 52 (0) | 15 (0) | 3.8±0.8 | 3.6±1.1 | 82±24 | 108±18<sup>a</sup> | I-V, VI-VII |
| Herbert et al. 2003 | 17 (0) | 15 (0) | 9.0±1.4<sup>a</sup> | 9.0±1.4<sup>a</sup> | 100±15<sup>c</sup> | 115±15<sup>b</sup> | Cb |
| Kaufmann et al. 2003 | 10 (0) | 22 (0) | 6.9±2.4 | 8.3±1.9 | 66±14 | 121±9 | Vermis, I-V, VI-VII, VIII-X |
| Pierce and Courchesne 2001 | 14 (2) | 14 (4) | 3.8±1.1 | 4.4±1.2 | 84±24 | 110±12 | I-V, VI-VII |
| Courchesne et al. 2001 | 60 (0) | 52 (0) | 6.2±3.5 | 8.1±3.5 | 79±26 | 115±15<sup>a</sup> | Cb, WM, GM |
| Hardan et al. 2001 | 22 (0)* | 22 (0)* | 22.4±10.1 | 22.4±10.0 | 100±15 | 100±14 | Vermis, I-V, VI-VII, VIII-X, Cb |
| Elia et al. 2000 | 22 (0) | 11 (0) | 10.9±4.0 | 10.9±2.9 | 55±10<sup>e</sup> | 115±15<sup>b</sup> | Vermis, VI-VII |
| Carper and Courchesne 2000 | 42 (0) | 29 (0) | 5.4±1.7 | 6.0±1.8 | 80±22 | 114±12 | VI-VII |
| Levitt et al. 1999 | 8 (-) | 21 (-) | 12.5±2.2 | 12.0±2.8 | 83±12 | 115±11 | VIII-X |
| Piven et al. 1997 | 35 (9) | 36 (16) | 18.0±4.5 | 20.2±3.8 | 91±20 | 102±13 | Cb |
| Ciesielski et al. 1997 | 9 (4) | 10 (3) | 16.8±5.2<sup>a</sup> | 16.6±5.4<sup>a</sup> | 93±13<sup>a</sup> | 119±10<sup>a</sup> | I-V, VI-VII |
| Hashimoto et al. 1995 | 96 (24) | 112 (47) | 6.4±4.9 | 7.2±5.2 | 60±25 | 99±18 | Vermis, I-V, VI-VII, VIII-X |
| Courchesne et al. 1994 | 50 (9) | 53 (10) | 16.5±11.6<sup>a</sup> | 18.8±10.4<sup>a</sup> | 81±31<sup>a</sup> | 115±15<sup>b</sup> | I-V, VI-VII |
| Piven et al. 1992 | 15 (0) | 15 (0) | 27.7±10.7 | 28.8±5.6 | 92±23 | 130±0 | I-V, VI-VII |
| Holttum et al. 1992 | 18 (0) | 18 (0) | 20.2±8.1 | 20.2±8.3 | 94±12 | 95±12 | Vermis, I-V, VI-VII, VIII-X |
| Garber and Ritvo 1992 | 12 (3) | 12 (4) | 27.2±5.3 | 26.4±3.6 | 95±15<sup>d</sup> | 115±15<sup>b</sup> | Vermis, I-V, VI-VII |
| Kleiman, Neff, and Rosman 1992 | 13 (3) | 17 (8) | 7.7±5.7 | 7.0±4.3<sup>a</sup> | 52±17 | 115±15<sup>b</sup> | I-V, VI-VII |
| Ritvo et al. 1988 | 15 (4) | 15 (4) | 11.6±4.1 | 11.6±4.1 | 75±15 | 115±15<sup>b</sup> | Vermis, I-V, VI-VII |

Volumes: Cb, cerebellum; WM, cerebellum white matter; GM, cerebellum grey matter
Mid-sagittal areas: whole vermis and its lobules numbered (I-V, anterior; VI-VII, superior-posterior; VIII-X, inferior-posterior).
ASD, autism spectrum disorders; Ctrl, normal controls; F, female. Age and IQ data: mean±SD
<sup>a</sup> Mean and/or standard deviation extrapolated from minimum and maximum
<sup>b</sup> IQ for normal control group supposed to be 115±15
<sup>c</sup> IQ for high functioning autism group supposed to be 100±15
<sup>d</sup> IQ for medium to high functioning autism group supposed to be 95±15
<sup>e</sup> IQ for low functioning autism group supposed to be 55±10
\* Cerebellar volume was estimated for only 16 ASD and 19 control subjects in Hardan et al. 2001

#### Mean effect size

Significant mean effects were found for 3 out of the 7 regions studied (whole cerebellum: p=0.049, white matter: p=0.011 and Lobules VI-VII: p=0.022, See **Table 2**). Compared with controls, individuals with ASD displayed larger cerebellar white matter volume (Cohen’s d=0.31, 95% CI: [0.07, 0.55]) and smaller areas for vermal VI-VII lobules (Cohen’s d=-0.24, 95% CI: [-0.44, -0.03]). The effect of diagnosis on whole cerebellar volume was barely statistically significant (Cohen’s d=0.23, 95% CI: [0.00, 0.45], see forest plot in **Figure 3**).

**Table 2.** Standardised mean difference: combination of effect sizes

| Region | Number of studies | Random Effects model |  |  | Meta-regression |  |  |
| --- | --- | --- | --- | --- | --- | --- | --- |
| | | Effect size (Cohen's d) | p-value | Heterogeneity ($I^2$) | Age impact (year<sup>-1</sup>) | IQ impact (IQ<sup>-1</sup>) | Residual heterogeneity ($I^2$) |
| Cerebellum | 16 | **0.23 (0.00, 0.45)** | **0.049** | 66% (42%, 80%) | -0.026 (-0.057, 0.005) | 0.014 (-0.002, 0.030) | **57% p=0.002** |
| Cerebellum WM | 6 | **0.31 (0.07, 0.55)** | **0.011** | 19% (0%, 64%) | **-0.153 (-0.287, -0.019)** | **0.042 (0.007, 0.077)** | 0% p=0.943 |
| Cerebellum GM | 6 | -0.22 (-0.47, 0.03) | 0.085 | 16% (0%, 79%) | **0.152 (0.020, 0.284)** | -0.033 (-0.067, 0.002) | 0% p=0.837 |
| Whole vermis | 11 | -0.09 (-0.30, 0.13) | 0.416 | 21% (0%, 60%) | 0.011 (-0.039, 0.062) | 0.009 (-0.010, 0.029) | 0% p=0.657 |
| Lobules I-V | 16 | 0.00 (-0.22, 0.23) | 0.971 | **54% (20%, 74%)** | -0.019 (-0.054, 0.016) | 0.020 (0.003, 0.038) | 29% p=0.061 |
| Lobules VI-VII | 18 | **-0.24 (-0.44, -0.03)** | **0.022** | **50% (14%, 71%)** | 0.009 (-0.024, 0.041) | 0.010 (-0.006, 0.026) | **36% p=0.039** |
| Lobules VIII-X | 9 | -0.20 (-0.50, 0.10) | 0.192 | **56% (8%, 79%)** | -0.043 (-0.167, 0.080) | 0.008 (-0.038, 0.055) | **65% p=0.013** |

Values in parentheses represent 95% confidence intervals; WM white matter; GM: grey matter.

**Figure 3.**
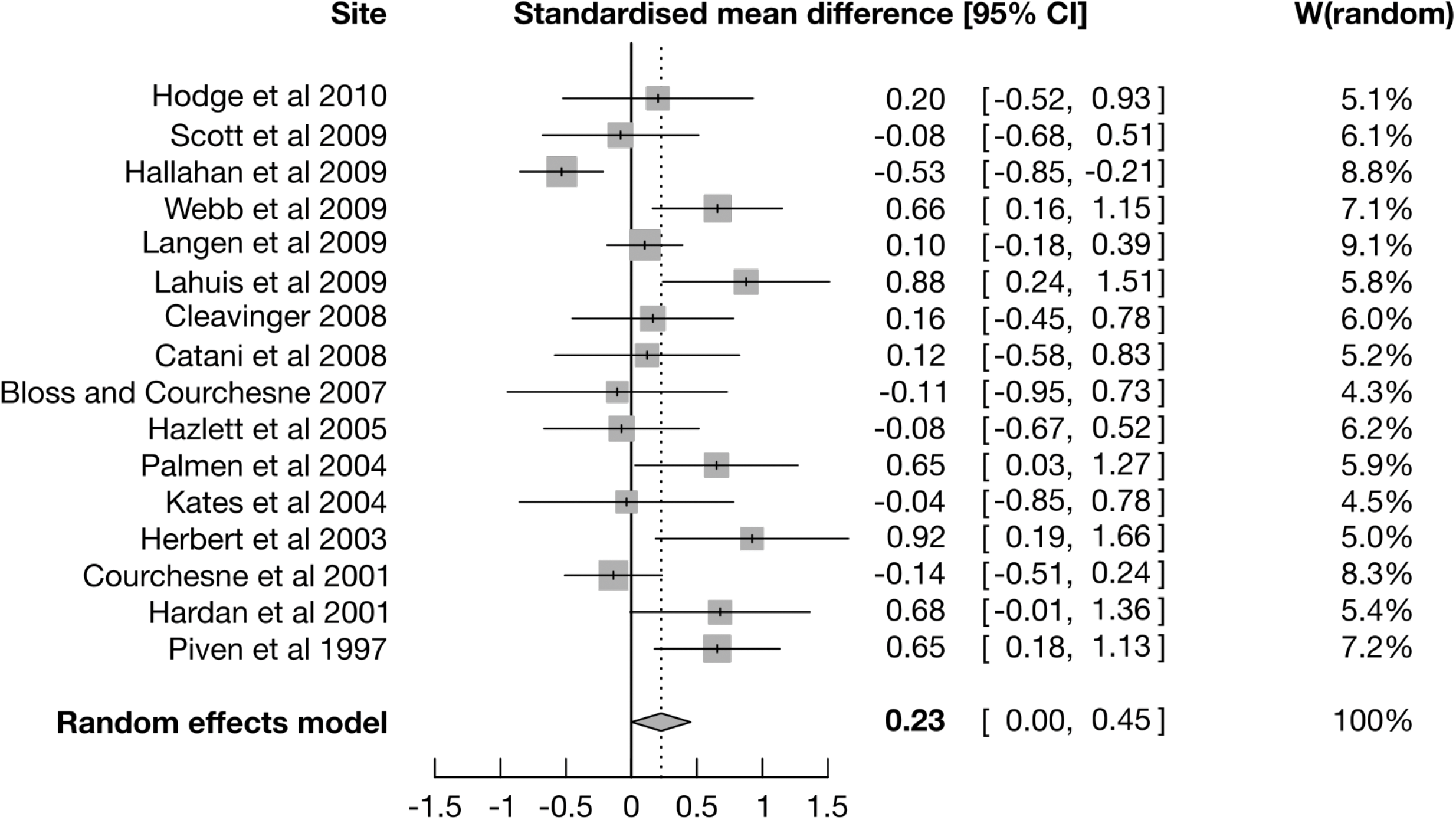
Forest plot for the cerebellum volume. For each study, the grey square is centred on the estimated standardised mean difference (SMD), the black segment illustrates the confidence interval (95%-CI), and the surface of the square is proportional to the number of subjects in the study. SMD is positive when the cerebellum volume is greater in ASD. W(random) represents the weight given to each study for the combination of effect sizes.

#### Heterogeneity and meta-regression

A statistically significant heterogeneity was found for 4 out of the 7 regions under study: cerebellum volume (p = 0.0001), vermal lobules I to V area (p = 0.0049), vermal lobules VI to VII area (p = 0.0081) and vermal lobules VIII to X areas (p = 0.019). Despite this high heterogeneity, our meta-regression did not show a significant impact of the age nor the IQ on total cerebellar volume. Age of individuals with ASD correlated reduced volume of cerebellar white matter (p = 0.025) and increased volume of cerebellar grey matter (p = 0.24) compared with controls. However, the number of studies included in the meta-regression may be insufficient to obtain reliable results. The IQ of individuals with ASD correlated with increased volume of the cerebellar white matter (p=0.0176) and increased area of vermal lobules I-V (p=0.019). Age and IQ did not seem to be the only factors producing heterogeneity: after the meta-regression, residual heterogeneity was still statistically significant for cerebellum volume (p=0.0020), vermal lobules VI-VII area (p=0.039) and vermal lobules VIII-X area (p=0.013). **Table 2** shows the results of the random effects models and meta-regressions for the cerebellum regions and **Figure 4** shows the observed effect sizes versus the expected effect from the meta-regression on whole cerebellum volume.

**Figure 4.**
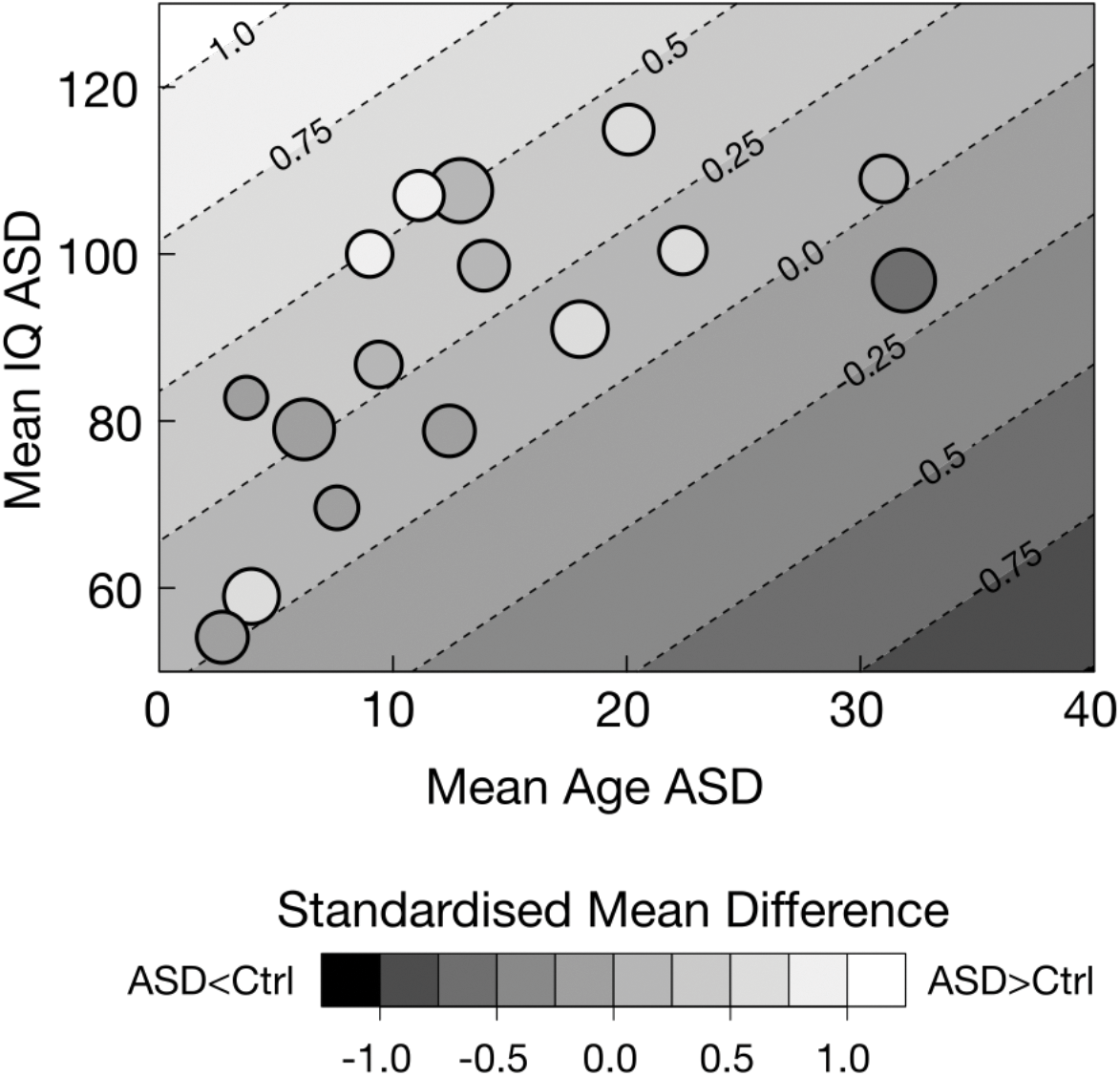
Meta-regression on standardised mean difference for the whole cerebellum volume.

In **Figure 4**, each study is represented by a circle positioned according to the mean age and the mean IQ of the ASD group. The surface of each circle is inversely proportional to estimated error variance (Thompson and Higgins 2002), and the grey level corresponds to the observed effect size (SMD, Standardised Mean Difference): light grey means a larger cerebellum volume on average for the ASD group compared to the control group, dark grey means a smaller volume. The background grey level gradient represents the effect size (of ASD versus control difference) predicted by the meta-regression. The grey levels of the circles were not well ordered, and they were often far from the background grey level, reflecting that age and IQ of ASD alone were not good predictors of mean cerebellar volume difference between ASD and controls.

#### Statistical power

Despite a small mean effect size estimated at Cohen’s d=0.23 for the whole cerebellum volume, 44% of the studies reported a significant result. If the actual effect size were fixed at this value, the mean statistical power for all the studies would be only 14%, that is, only 14% chances of detecting such a small effect size. The heterogeneity in age and IQ across studies appeared to limit the statistical power: mean achieved statistical power increased to 20% when taking into account the variations induced by age and IQ estimated by the meta-regression (the value is, however, still much lower than the observed 44% rate of detection which agrees with the fact that much of the heterogeneity was not explained).

#### Publication bias and p-hacking

Egger’s test reported a statistically significant funnel plot asymmetry only for the whole vermis area (p=0.002). None of the remaining regions presented a significant funnel plot asymmetry (**Figure 5**).

**Figure 5.**
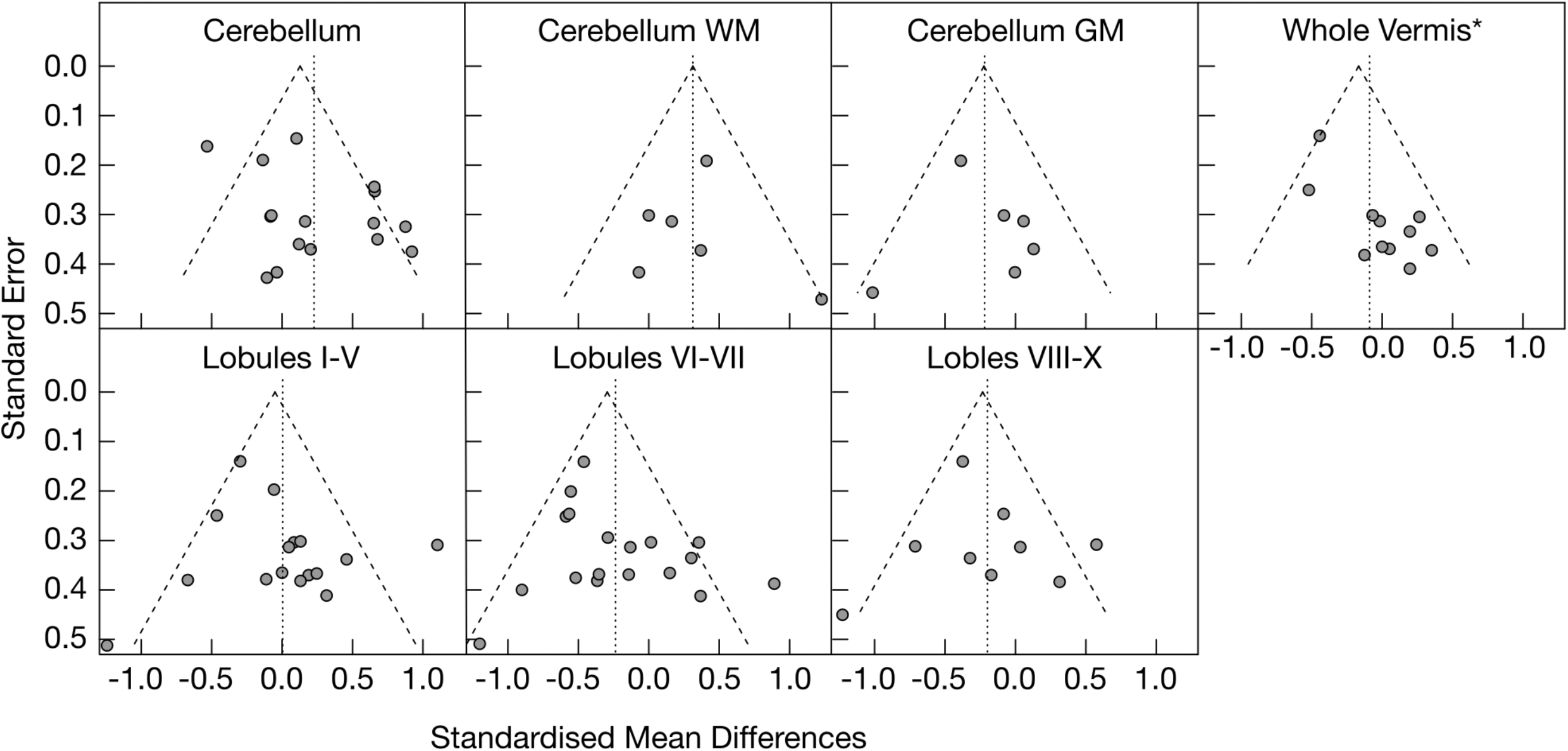
Funnel plot for the different regions WM: white matter; GM: grey matter; * statistically significant asymmetry of the funnel plot (p=0.002). The dotted line represents the meta-analytic effect size whereas the two dashed lines represent the boundaries of the 95% confidence interval for effect combined with the fixed effect model. Heterogeneity: I^2^=65.9%, *τ*^2^=0.1235, p=0.0001.

Our analyses of the p-curves were not conclusive on the presence of p-hacking. In 5 out of the 7 regions there were only 2 or 3 p-values <0.05, which resulted in very imprecise estimations with wide confidence intervals. Total cerebellum and lobules VI-VII had each 7 p-values <0.05, which still produced unreliable estimations. P-curve analyses suggested low statistical power for both regions (Cerebellum: 22%, 95% CI [5%, 67%] and Lobules VI-VII: 38%, 95% CI [6%, 78%]). The statistical powers inferred from the p-curves were not incompatible with the respective rates of significant studies, but a gain, confidence intervals were very wide. **Table 3** summarises the results for the different publication bias and p-hacking analyses. **Figure 6** shows the observed p-curve for the total cerebellar volume.

**Table 3.** Standardized Mean Difference: publication bias and p-hacking

| Region | Rate of significant studies | Mean achieved power | Egger's test | P-curve inferred power |
| --- | --- | --- | --- | --- |
| Cerebellum | 44% (7/16) | 20% | $p = 0.064$ | 22% (5%, 67%) |
| Cerebellum WM | 33% (2/6) | 28% | $p = 0.870$ | 19% (5%, 90%) |
| Cerebellum GM | 33% (2/6) | 22% | $p = 0.784$ | 5% (5%, 59%) |
| Whole vermis | 18% (2/11) | 16% | **$p = 0.002$** | 32% (5%, 93%) |
| Lobules I-V | 19% (3/16) | 13% | $p = 0.353$ | 55% (5%, 94%) |
| Lobules VI-VII | 39% (7/18) | 19% | $p = 0.165$ | **38% (6%, 78%)** |
| Lobules VIII-X | 33% (3/9) | 13% | $p = 0.726$ | 47% (5%, 92%) |

Mean achieved powers are based on the effect sizes estimated by the meta-regression (taking into account the impacts of average age and IQ of ASD groups)
Values in parentheses for p-curve inferred powers are 90% confidence intervals

**Figure 6.**
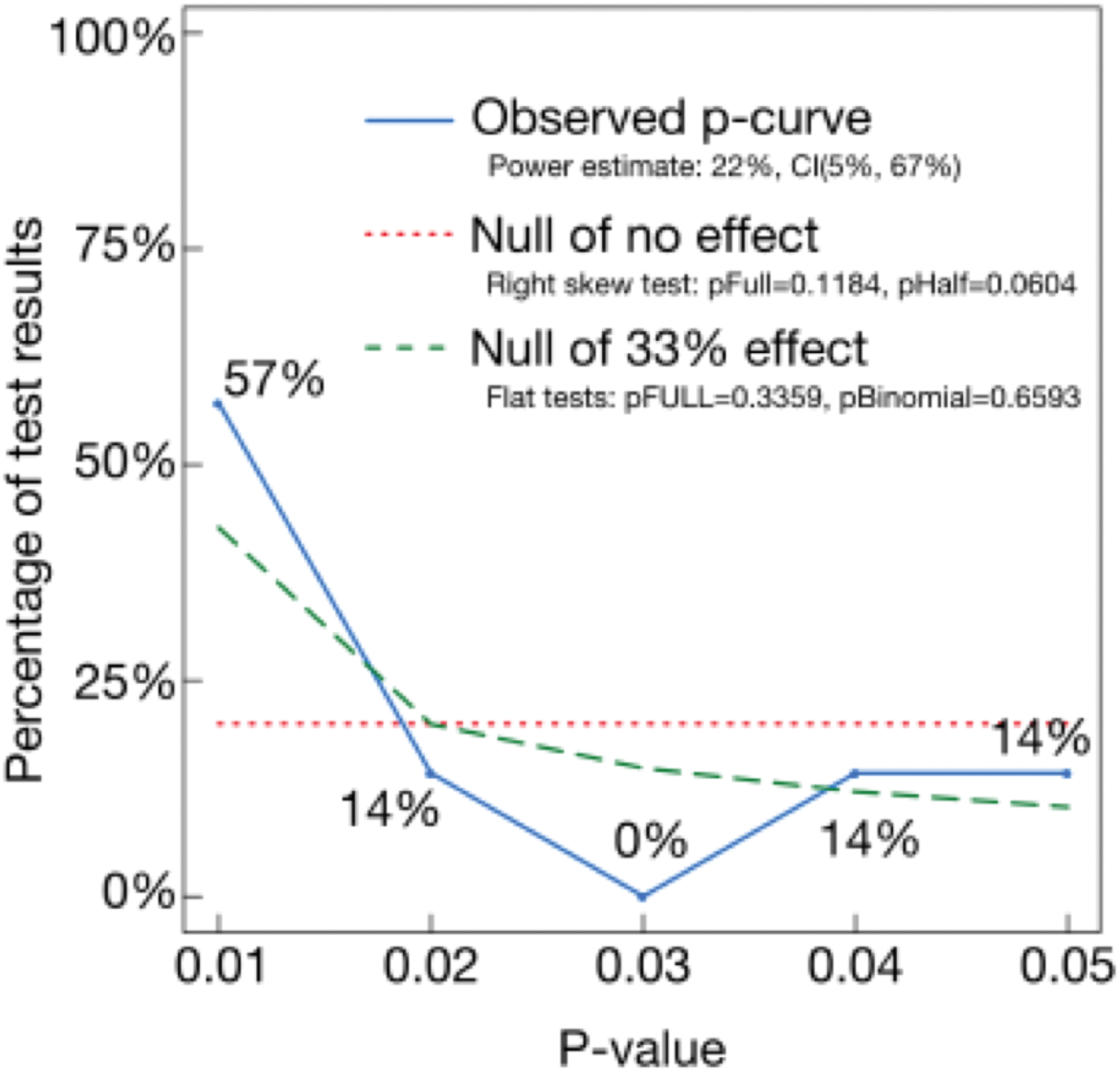
P-curve of studies with a significant result for the cerebellum volume. Null of: expected p-curve in case of. Note: The observed p-curve includes 7 studies with statistically significant results (p<.05). Among them, 5 had p<.025. There were 8 additional studies excluded from the p-curve analysis because their results did not pass significance (p>.05).

#### Meta-analysis of variability (log-variance ratio)

A barely statistically significant (uncorrected) effect was found for vermal lobules VIII-X areas, suggesting larger volume variations for the ASD groups (p=0.049). No significant effect was found for any other regions. Heterogeneity was statistically significant only for the whole cerebellar volume (p = 0.0052, uncorrected). As for the meta-analysis of mean differences, taking into account age and IQ in the meta-regression did not significantly reduce the heterogeneity (see **Table 4**).

**Table 4.** Log-variance ratio: combination of effect sizes

| Region | Number of studies | Random Effects model |  |  | Meta-regression |  |  |
| --- | --- | --- | --- | --- | --- | --- | --- |
|  |  | Effect size | p-value | I<sup>2</sup> indicator of heterogeneity | Age impact (year<sup>-1</sup>) | IQ impact (IQ<sup>-1</sup>) | I<sup>2</sup> indicator of residual heterogeneity |
| Cerebellum | 16 | 0.02 (-0.27, 0.31) | 0.898 | **54% (19%, 74%)** | 0.018 (-0.021, 0.057) | -0.014 (-0.034, 0.006) | **45% p = 0.033** |
| Cerebellum WM | 6 | -0.20 (-0.56, 0.15) | 0.259 | 0% (0%, 37%) | -0.039 (-0.235, 0.157) | 0.014 (-0.037, 0.066) | 0% p = 0.640 |
| Cerebellum GM | 6 | 0.08 (-0.43, 0.60) | 0.749 | 44% (0%, 78%) | **0.254 (0.058, 0.450)** | **-0.064 (-0.115, -0.012)** | 0% p = 0.530 |
| Whole vermis | 11 | 0.03 (-0.21, 0.28) | 0.778 | 0% (0%, 39%) | -0.013 (-0.089, 0.063) | 0.001 (-0.028, 0.031) | 0% p = 0.621 |
| Lobules I-V | 15* | 0.01 (-0.25, 0.26) | 0.962 | 17% (0%, 54%) | 0.014 (-0.039, 0.067) | 0.001 (-0.024, 0.027) | 26% p = 0.185 |
| Lobules VI-VII | 18 | 0.10 (-0.09, 0.29) | 0.311 | 0% (0%, 49%) | **0.040 (0.001, 0.079)** | -0.008 (-0.026, 0.010) | 0% p = 0.658 |
| Lobules VIII-X | 9 | 0.25 (0.00, 0.51) | **0.049** | 0% (0%, 32%) | 0.034 (-0.077, 0.145) | -0.009 (-0.048, 0.030) | 0% p = 0.717 |

\* An article (Ciesielski et al. 1997) had to be excluded from the meta-analysis on the log-variance ratio for lobules I-V because the standard deviation in each group was not available, and the one reconstructed from the reported Student t value (for the standardised Mean Difference analyses) did not allow us to infer a different value for each group.

### 2. Analysis of the ABIDE data

From the original 1112 subjects, 328 patients (61% of the evaluated subjects) and 353 controls (62% of the evaluated subjects) were retained for further analysis after quality control (see **Table 5** for a description of selected subjects by ABIDE site). Among the excluded subjects, 411 subjects did not pass the quality control step and 20 subjects were excluded because of unavailable FIQ (Full-scale Intelligence Quotient).

**Table 5.** Demographics of ABIDE subjects for analysis

| Site | Institution | N<sub>ASD</sub> (F)* | N<sub>Ctrl</sub> (F)* | Age<sub>ASD</sub> | Age<sub>Ctrl</sub> | IQ<sub>ASD</sub> | IQ<sub>Ctrl</sub> |
| --- | --- | --- | --- | --- | --- | --- | --- |
| Caltech | California Institute of Technology, USA | 14/19 (3/4) | 14/19 (4/4) | 27.4±10.7 | 30.1±12.2 | 108±13 | 112±9 |
| CMU | Carnegie Mellon University, USA | 0/14 (0/3) | 0/13 (0/3) | - | - | - | - |
| KKI | Kennedy Krieger Institute, USA | 17/22 (3/4) | 29/33 (9/9) | 10.1±1.5 | 10.1±1.3 | 95±16 | 114±9 |
| Leuven | University of Leuven, Belgium | 29/29 (3/3) | 32/35 (5/5) | 17.8± 5.0 | 17.8±5.0 | 101±16 | 113±11 |
| MaxMun | Ludwig Maximilian University Munich, Germany | 14/24 (2/3) | 18/33 (4/4) | 23.8± 14.0 | 27.0±10.3 | 108±15 | 109±12 |
| NYU | New York University Langone Medical Center, USA | 29/79 (5/11) | 53/105 (17/26) | 13.3± 5.3 | 17.1±6.7 | 111±17 | 114±11 |
| OHSU | Oregon Health and Science University, USA | 11/13 (0/0) | 15/15 (0/0) | 11.1± 1.9 | 10.1±1.1 | 106±22 | 116±11 |
| Olin | Olin, Institute of Living at Hartford Hospital, USA | 15/20 (2/3) | 16/16 (2/2) | 16.5± 3.0 | 16.9±3.7 | 110±18 | 115±17 |
| Pitt | University of Pittsburgh, School of Medicine, USA | 23/30 (3/4) | 20/27 (4/4) | 19.4± 7.9 | 17.4±5.1 | 110±15 | 110±10 |
| SBL | Netherlands Institute for Neurosciences, Netherlands | 0/15 (0/0) | 0/15 (0/0) | - | - | - | - |
| SDSU | San Diego State University, USA | 7/14 (1/1) | 17/22 (5/6) | 14.8±1.9 | 14.1±2.1 | 111±19 | 109±11 |
| Stanford | Stanford University, USA | 5/20 (2/4) | 5/20 (2/4) | 10.0±1.6 | 11.0±2.0 | 108±26 | 121±9 |
| Trinity | Trinity Centre for Health Sciences, Ireland | 22/24 (0/0) | 23/25 (0/0) | 17.7±3.4 | 17.2±3.8 | 109±16 | 112±12 |
| UCLA | University of California, Los Angeles, USA | 49/62 (6/7) | 43/47 (6/6) | 13.1±2.4 | 13.0±2.0 | 101±13 | 107±11 |
| UM | University of Michigan, USA | 16/68 (2/10) | 6/77 (2/18) | 13.5±2.5 | 16.9±1.1 | 107±20 | 114±5 |
| USM | University of Utah, School of Medicine, USA | 52/58 (0/0) | 38/43 (0/0) | 22.6±7.9 | 21.8±7.9 | 100±16 | 115±13 |
| Yale | Yale Child Study Center, USA | 25/28 (8/8) | 24/28 (8/8) | 13.0±3.0 | 12.8±2.8 | 94±21 | 106±18 |
|  | Total | 328/539 (40/65) | 353/573 (68/99) | 16.8±7.5 | 16.8±7.4 | 104±17 | 112±12 |

ASD, autism spectrum disorders; Ctrl, controls; F, female. Age and IQ data: mean±SD
\* Number of subjects retained / Number of subjects segmented (Number of female subjects retained / Number of female subjects segmented)

#### Allometry

The scaling between cerebellar volume and brain volume was not isometric (i.e., the volume of the cerebellum was not directly proportional to brain volume). The scaling factor was estimated at 0.518 (95% CI: 0.457, 0.578), statistically significantly smaller than 1 (which would be the case for isometry) (**Figure 7**). Our estimation shows that large brains have a proportionally smaller cerebellum. Because of this non-proportional relationship, the normalisation of cerebellar volumes by total brain volume should not be used to control for group differences. This validates our choice of using instead brain volume as covariate in our linear models.

**Figure 7.**
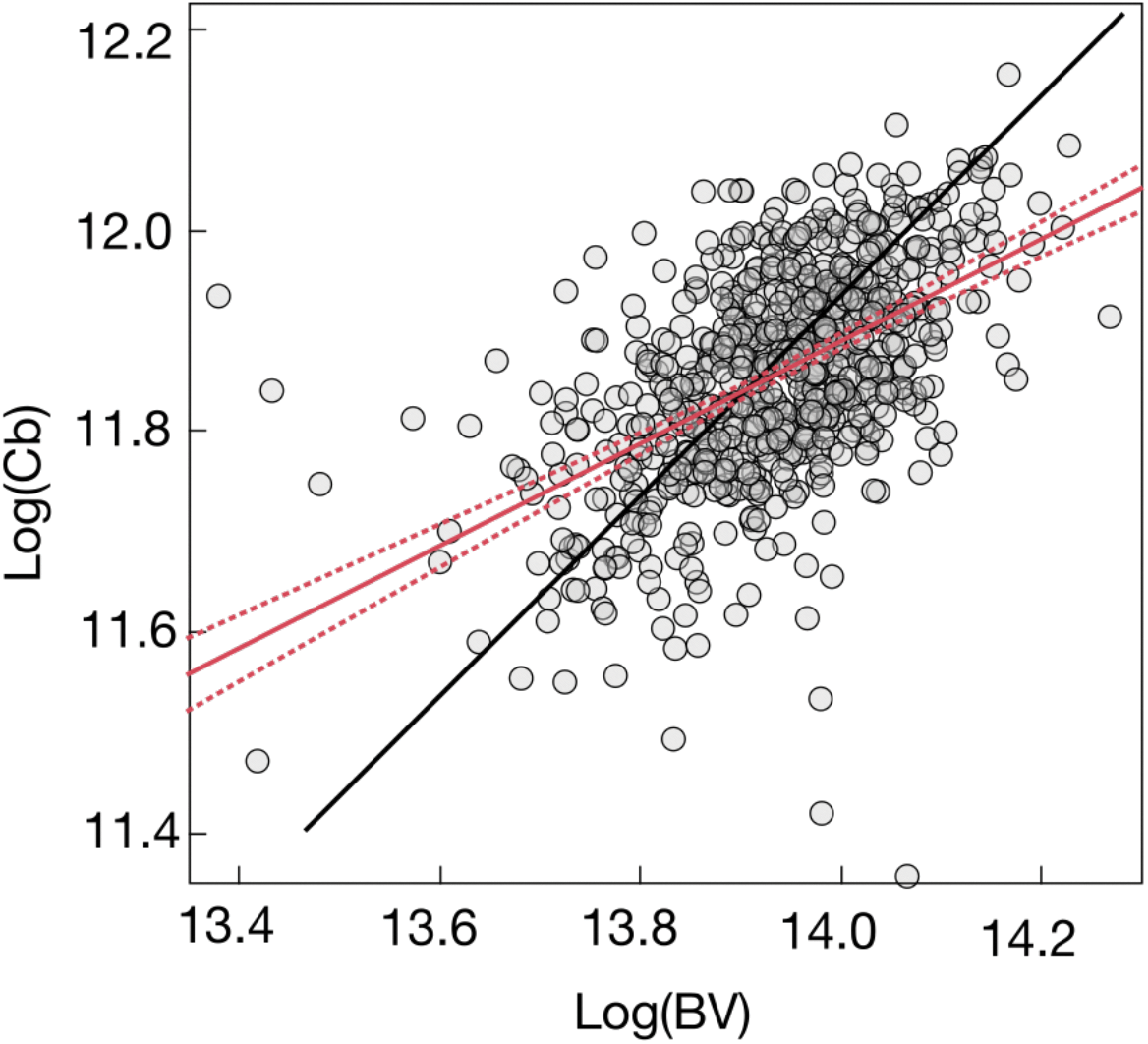
Log of cerebellar volume versus log of total brain volume.

#### Linear model

A very significant site effect was found for each volume, this effect was also different between patients and controls for the whole cerebellum volume and the cerebellum grey matter volume. The effect of diagnosis group for the total cerebellum volume was not significant (estimated at -0.59 cm3, Cohen’s d=-0.04, 95% CI: [-0.16, 0.09], in the first model – including group, age, IQ, scanning site, sex and brain volume as fixed effects; and at -1.22 cm3, Cohen’s d=-0.08, 95% CI: [-0.27, 0.11], in the second model – including in addition the interactions of group with age, IQ, sex, brain volume and scanning site). These estimations were more precise than in the meta-analysis of the literature, with narrower confidence intervals. Despite this, we did not find any significant difference between patients and controls. We did not find either any impact of age, sex, IQ or brain volume on cerebellar volume that would differ by diagnosis group.

The analysis of cerebellar subregions (white and grey matter volumes) did not lead to a statistically significant group effect either. See **Table 6** for the results of the linear model with group as main effect and **Table 7** for the results model including the interaction of group with the other variables.

**Table 6.** Effects of group, site, age, IQ, sex and BV on cerebellar volumes

|  | Whole cerebellum volume | Cerebellum white matter volume | Cerebellum grey matter volume |
| --- | --- | --- | --- |
| Mean volume ± SD ($\text{cm}^3$) | 141.7 ± 15.0 | 30.2 ± 4.2 | 111.5 ± 12.6 |
| Group (ASD) effect ($\text{cm}^3$) | -0.59 (-2.46, 1.28)<br>p = 0.5373 | -0.30 (-0.80, 0.21)<br>p = 0.2517 | -0.29 (-1.88, 1.29)<br>p = 0.7171 |
| Site effect ($\text{cm}^3$) | from -7.48 to 11.08<br>p < 0.0001 | from -3.07 to 8.53<br>p < 0.0001 | from -7.61 to 13.93<br>p < 0.0001 |
| Age effect ($\text{cm}^3/\text{year}$) | -0.34 (-0.50, -0.19)<br>p < 0.0001 | 0.12 (0.08, 0.16)<br>p < 0.0001 | -0.46 (-0.60, -0.33)<br>p < 0.0001 |
| IQ effect ($\text{cm}^3/\text{IQ}$) | 0.05 (-0.01, 0.11)<br>p = 0.1094 | -0.01 (-0.03, 0.00)<br>p = 0.1106 | 0.06 (0.01, 0.12)<br>p = 0.0167 |
| Sex (F) effect ($\text{cm}^3$) | -4.71 (-7.38, -2.04)<br>p = 0.0006 | 0.52 (-0.21, 1.24)<br>p = 0.1613 | -5.22 (-7.49, -2.96)<br>p < 0.0001 |
| BV effect | 0.061 (0.053, 0.070)<br>p < 0.0001 | 0.018 (0.016, 0.021)<br>p < 0.0001 | 0.043 (0.036, 0.050)<br>p < 0.0001 |

SD, standard deviation; ASD, autism spectrum disorders; IQ, intelligence quotient; M, male; BV, brain volume
The effect for a continuous factor (age or IQ) represents the mean variation induced by the variation of the factor.
The effect for a binary factor (group or sex) represents the mean variation induced by being in the group in brackets from the other group.
The effect for the site represents the mean variation induced by being in a site from the mean of the sites.
The values in brackets represent the 95% confidence intervals for the estimated effects.

**Table 7.** Effects of group combined with site, age, IQ, sex and BV on cerebellar volumes

|  | Whole cerebellum volume | Cerebellum white matter volume | Cerebellum grey matter volume |
| --- | --- | --- | --- |
| Mean volume ± SD ($\text{cm}^3$) | 141.7 ± 15.0 | 30.2 ± 4.2 | 111.5 ± 12.6 |
| Group (ASD) effect ($\text{cm}^3$) | -1.22 (-4.12, 1.68)<br>p = 0.4100 | -0.67 (-1.46, 0.12)<br>p = 0.0969 | -0.55 (-3.01, 1.91)<br>p = 0.6616 |
| Site effect ($\text{cm}^3$) | from -7.77 to 10.97<br>p < 0.0001 | from -3.11 to 8.52<br>p < 0.0001 | from -7.83 to 13.85<br>p < 0.0001 |
| Site * Group (ASD) effect ($\text{cm}^3$) | from -16.70 to 10.58<br>p = 0.0082 | from -2.99 to 2.34<br>p = 0.2277 | from -13.71 to 8.96<br>p = 0.0125 |
| Age effect ($\text{cm}^3/\text{year}$) | -0.35 (-0.51, -0.19)<br>p < 0.0001 | 0.12 (0.08, 0.17)<br>p < 0.0001 | -0.47 (-0.61, -0.34)<br>p < 0.0001 |
| Age * Group (ASD) effect ($\text{cm}^3/\text{year}$) | -0.08 (-0.40, 0.23)<br>p = 0.5989 | -0.01 (-0.10, 0.07)<br>p = 0.7863 | -0.07 (-0.34, 0.19)<br>p = 0.5942 |
| IQ effect ($\text{cm}^3/\text{IQ}$) | 0.05 (-0.01, 0.12)<br>p = 0.1261 | -0.01 (-0.03, 0.01)<br>p = 0.1629 | 0.07 (0.01, 0.12)<br>p = 0.0246 |
| IQ * Group (ASD) effect ($\text{cm}^3/\text{IQ}$) | 0.00 (-0.13, 0.14)<br>p = 0.9436 | -0.00 (-0.04, 0.04)<br>p = 0.9954 | 0.00 (-0.11, 0.12)<br>p = 0.9321 |
| Sex (F) effect ($\text{cm}^3$) | -4.62 (-7.34, -1.89)<br>p = 0.0009 | 0.46 (-0.29, 1.20)<br>p = 0.2280 | -5.07 (-7.39, -2.76)<br>p < 0.0001 |
| Sex (F) * Group (ASD) effect ($\text{cm}^3$) | -0.10 (-1.46, 1.27)<br>p = 0.8904 | -0.86 (-2.35, 0.62)<br>p = 0.2539 | 1.25 (-3.38, 5.87)<br>p = 0.5970 |
| BV effect | 0.062 (0.053, 0.070)<br>p < 0.0001 | 0.019 (0.016, 0.021)<br>p < 0.0001 | 0.043 (0.036, 0.050)<br>p < 0.0001 |
| BV * Group (ASD) effect | -0.003 (-0.020, 0.015)<br>p = 0.7681 | -0.000 (-0.005, 0.004)<br>p = 0.8367 | -0.002 (-0.017, 0.013)<br>p = 0.7784 |

SD, standard deviation; ASD, autism spectrum disorders; IQ, intelligence quotient; M, male; BV, brain volume
The effect for a continuous factor (age, IQ or BV) represents the mean variation induced by the variation of the factor.
The effect for a binary factor (group or sex) represents the mean variation induced by being in the group in brackets from the other group.
The effect for the site represents the mean variation induced by being in a site from the mean of the sites.
The values in brackets represent the 95% confidence intervals for the estimated effects.

#### ABIDE meta-analysis

The results of the analyses using the meta-analytical approach agreed with those of the direct fit of linear models. Despite a smaller number of subjects than in the meta-analysis of the literature (**Table 8**), the confidence intervals for the combined mean differences were narrower. This was due to a smaller estimated between-study variance among ABIDE sites compared with that among articles in the literature. We did not find a statistically significant standard mean difference between patients and controls: d=0.06 (95 % CI: -0.12; 0.24) (**Figure 8**). In the meta-regression, no significant effect was found for age (p=0.888) nor IQ (p=0.726). Table 9 describes the characteristics of the subjects selected for the preservation of age and sex matching between patients and controls. **Figure 8** shows the forest plot of ABIDE sites combined with the random effects model. The meta-analytical approach did not show either any statistically significant difference in the variability of volume measures between patients and controls (log-variance ratio=0.17, 95 % CI: -0.18; 0.52). **Table 9** summarises the results of the ABIDE meta-analysis on standardised mean difference and **Table 10** summarises the results of the ABIDE meta-analysis on log-variance ratio.

**Table 8.** ABIDE sites included in the meta-analysis

| Site | N<sub>ASD</sub> (F) | N<sub>Ctrl</sub> (F) | Age<sub>ASD</sub> | Age<sub>Ctrl</sub> | IQ<sub>ASD</sub> | IQ<sub>Ctrl</sub> |
| --- | --- | --- | --- | --- | --- | --- |
| CALTECH | 14 (3) | 14 (4) | 27.4±10.7 | 30.1±12.2 | 108.0±13.4 | 111.9±8.7 |
| KKI | 17 (3) | 29 (9) | 10.1±1.5 | 10.1±1.3 | 95.5±15.7 | 114.4±9.0 |
| LEUVEN | 29 (3) | 32 (5) | 17.8±5.0 | 17.8±5.0 | 101.4±16.4 | 112.6±11.2 |
| MAX | 12 (0) | 14 (0) | 21.6±13.5 | 24.9±10.1 | 105.5±14.9 | 108.9±12.6 |
| NYU | 24 (4) | 50 (16) | 13.8±5.5 | 16.4±6.2 | 109.5±15.6 | 114.5±11.5 |
| OHSU | 9 (0) | 15 (0) | 10.5±1.5 | 10.1±1.1 | 101.3±21.7 | 115.7±11.1 |
| OLIN | 13 (0) | 14 (0) | 16.5±3.1 | 16.9±3.8 | 108.4±18.2 | 115.1±16.9 |
| PITT | 21 (3) | 20 (4) | 17.9±6.6 | 17.4±5.1 | 110.4±13.9 | 110.2±10.4 |
| SDSU | 6 (0) | 12 (0) | 15.1±1.7 | 14.5±1.6 | 114.3±19.3 | 112.5±9.5 |
| TRINITY | 22 (0) | 23 (0) | 17.7±3.4 | 17.2±3.8 | 109.2±15.6 | 111.9±12.2 |
| UCLA | 49 (6) | 43 (6) | 13.1±2.4 | 13.0±2.0 | 101.5±13.1 | 106.6±10.9 |
| USM | 52 (0) | 38 (0) | 22.6±7.9 | 21.8±7.9 | 99.6±15.6 | 114.9±13.1 |
| YALE | 25 (8) | 24 (8) | 13.0±3.0 | 12.8±2.8 | 93.7±21.3 | 106.2±17.7 |
| Total | 293 (30) | 328 (52) | 17.0±7.4 | 16.6±7.2 | 103.0±16.6 | 111.9±12.3 |

**Figure 8.**
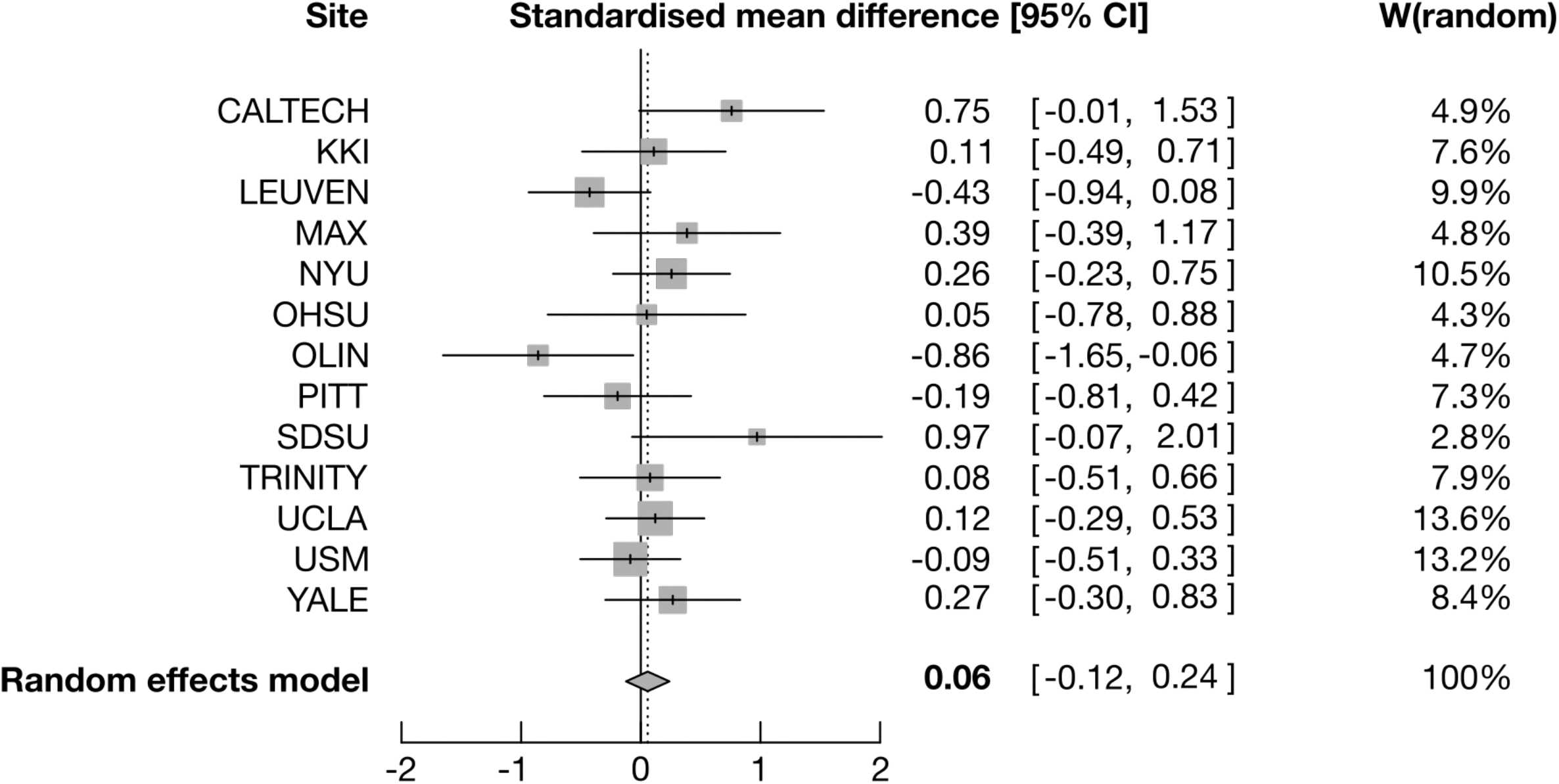
Forest plot for standardized mean difference on the cerebellar volume involving the different ABIDE sites. For each site, the grey square is centred on the estimated standardized mean difference (SMD), the black segment illustrates the confidence interval (95%-CI), and the surface of the square is proportional to the number of subjects in the site. SMD is positive when the cerebellum volume is greater in ASD. W(random) represents the weight given to each site for the combination of effect sizes. Heterogeneity: I^2^=32.6%, *τ*^2^=0.0192, p=0.1219.

**Table 9.**
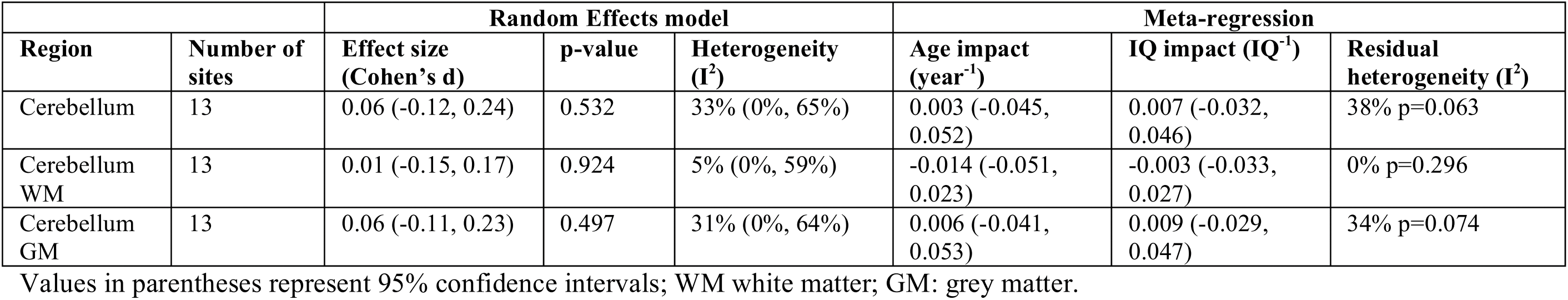
Standardised Mean Difference: combination of effect sizes from ABIDE site.

**Table 10.**
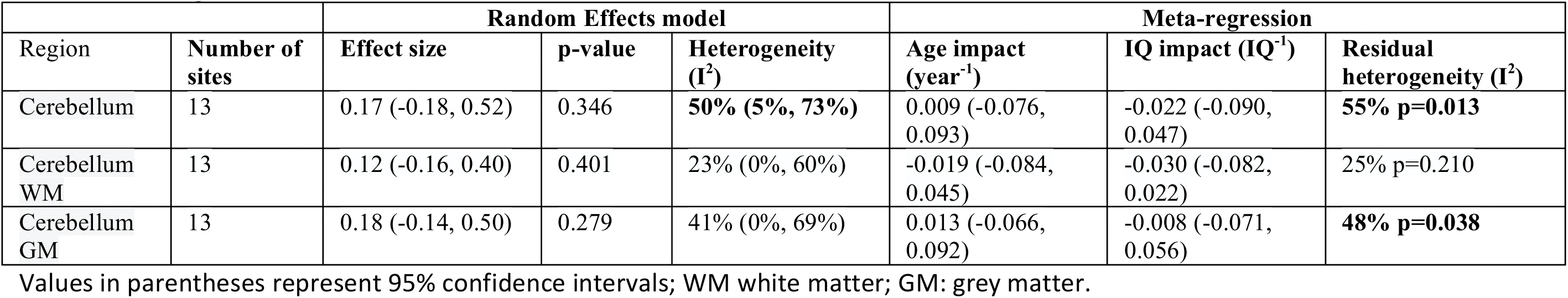
Log-variance ratio: combination of effect sizes form ABIDE sites

## Discussion

Neuroanatomical diversity appears to capture a substantial proportion of the risk to ASD (Sabuncu et al. 2016). However, and even though several candidate neuroanatomical biomarkers have been proposed, it is not yet clear which are exactly the neuroanatomical traits that more strongly influence diagnosis. In this report we looked at one specific structure, the cerebellum, that has been widely discussed in the literature. We performed a meta-analysis of the literature and an analysis of the data from the ABIDE project.

The meta-analysis of the literature did not show conclusive evidence for a difference between individuals with ASD and controls neither for the total cerebellar volume nor for its subregions. Whole cerebellum and cerebellar white matter volumes appeared slightly larger in ASD whereas vermal lobules VI-VII area was found to be slightly smaller in ASD, but the significance of these results would not survive correction for multiple comparisons. Compared with a previous meta-analysis on whole cerebellum volume and vermal areas (Stanfield et al. 2008), the effect sizes we computed were all smaller in absolute value. Specifically, Stanfield et al. (2008) reported a significant difference in the volume of the vermal lobules VIII-X. In our analysis, despite the fact that a larger number of studies was taken into account, we did not replicate their finding.

The combined effect sizes in the meta-analysis of the literature were small in general. The number of articles that reported statistically significant results was larger than expected given the mean achieved power. We investigated whether this excess could be due to publication bias or p-hacking. Publication bias occurs when studies with statistically significant results have higher chances to be published than those without. This is more likely to happen when the finding reported is central to the hypothesis made by an article. However, the main focus of many of the articles that we meta-analysed was not cerebellar volume, which should decrease the likelihood of bias. We studied publication bias by analysing the asymmetry of the funnel plot using Egger’s test. This type of analysis is not very sensitive and is able to detect only strong publication bias. We observed statistically significant funnel plot asymmetry only for the vermis area. Whereas this result suggests the presence of publication bias, it could also be due to greater inter-study heterogeneity in this specific measurement (where a few large studies reported negative effect sizes, whereas the majority of the other studies reported positive effect sizes)

We aimed at testing for p-hacking by analysing the p-curves: the distribution of p-values smaller than 0.05. The numbers of significant p-values reported, however, were not sufficient to draw a definite conclusion. In the case of the total cerebellar volume, where 7 out of 16 articles reported statistically significant findings, p-curve analysis did not reveal evidence for p-hacking. Overall, the analysis of the literature alone does not allow us to provide a definitive explanation for the excess of statistically significant findings.

One result that appeared very clearly was the strong heterogeneity of the findings in the literature. A certain degree of variability in the estimations of volume is expected, especially with the small sample sizes used. Heterogeneity tests aim at detecting a degree of variability that would go beyond this expectation. We found statistically significant heterogeneity for the estimations of 4 out of the 7 meta-analysed regions (i.e., whole cerebellar volume, lob I-V, lob VI-VII and lob VIII-IX, see table 2). Heterogeneity may be due to a combination of technical and physiological causes, for example, differences in MRI equipment, acquisition sequences, segmentation protocol, but also the age of the subjects, IQ distribution, etc. We analysed the effect of 2 of these factors, age and IQ, using meta-regression. First, our analysis did not reveal a differential effect of age or IQ on cerebellar volume for patients and controls. Second, residual heterogeneity was still statistically significant for 3 out of the 7 regions studied (whole cerebellar volume, lob VI-VII, lob VIII-IX). This indicates that sources other than age and IQ level may be causing significant heterogeneity in the literature.

The ABIDE project data provides a very interesting point of comparison for previous findings in the literature. The subjects in ABIDE come in most cases from research projects that had already been published, and should be then of similar characteristics as those in our literature meta-analysis. However, because the raw MRI and behavioural data is available, there is no issue of publication bias or p-hacking having an effect on the cohort selection. Additionally, the availability of raw data makes it possible to run methodologically homogenous analyses: same segmentation protocol, quality control procedures, and statistical analyses. There still remain, of course, many additional sources of heterogeneity due to the grassroots nature of the project (some of them being currently addressed in ABIDE II through an important harmonisation effort). Overall, the analyses of ABIDE data should provide a more precise, less heterogeneous estimation. The availability of raw data makes it also possible to engage a community effort to assess the impact of different methodological choices on the very same dataset.

We analysed all 1112 subjects from ABIDE using validated automatic computational neuroanatomy tools. After quality control and additional inclusion criteria we retained a group of 681 subjects. We had 85% power to detect the Cohen’s d=0.23 effect obtained from our meta-analysis of the literature (2-sided t-test, alpha=0.05). We did not find any statistically significant result, neither for mean total volumes, interaction with age or IQ. Our statistical analysis here was a linear model including group, age, sex, IQ and site as main effects, or additionally the interaction between group and the other covariates. Although the absence of group effect was clear, we repeated our analyses using the same meta-analytical procedure used to study the literature to rule out an eventual methodological artefact. Every ABIDE site was considered as a different source of data and we computed a meta-analytical effect size using a random effects model weighting of each site’s estimations by the inverse of the variance (thus giving more weight to sites with larger sample sizes, as in the case of the literature meta-analysis). Our meta-analysis was in agreement with the results of our linear models: a clear absence of group differences.

We can also use the ABIDE results to better understand publication bias in the literature. **Figure 9** shows a funnel plot combining the results for total cerebellar volume in the literature and the different ABIDE centres. Egger’s test for funnel plot asymmetry provides again a suggestion of publication bias (p=0.0478). More research, probably expanding the scope of our analyses to encompass more brain regions, would be required to better understand the underlying reasons of the excess of significant reports in the literature.

**Figure 9.**
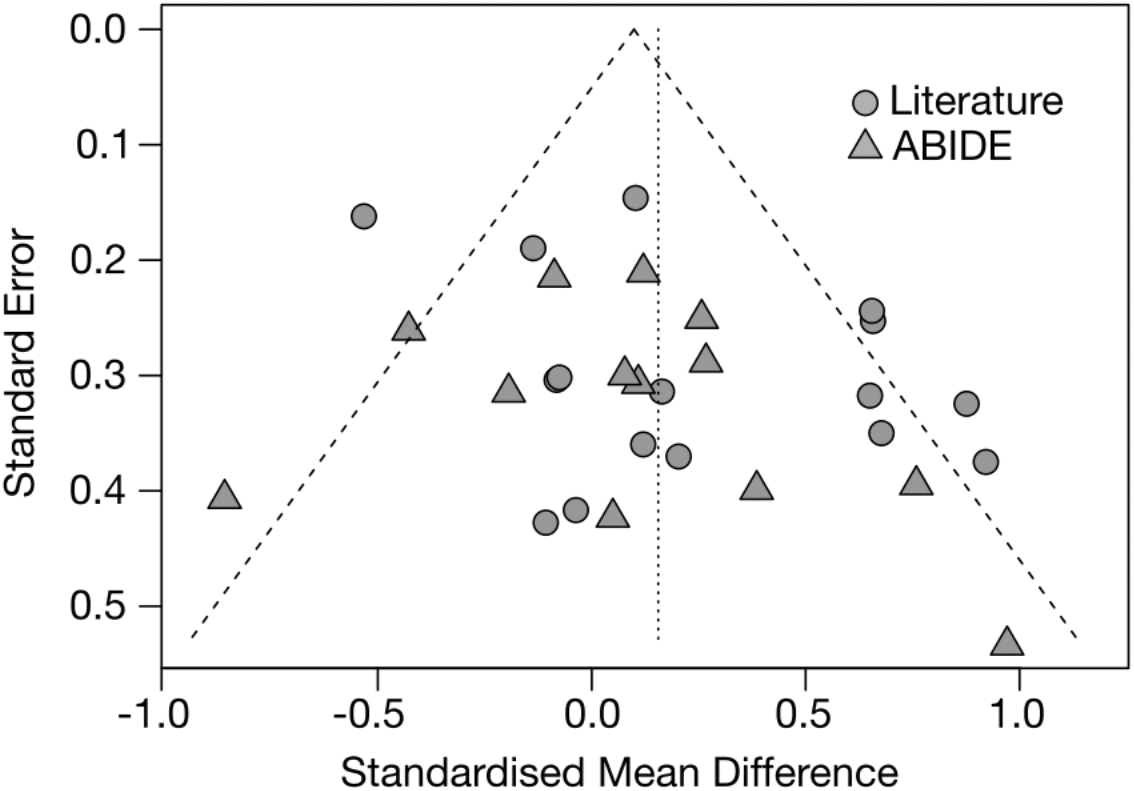
Funnel plot for the total cerebellum volume combining articles of the literature and ABIDE sites.

Interestingly, and although the site effect was substantial, the estimations from the ABIDE data were more precise than those from the literature (tighter confidence intervals), less heterogeneous. One reason for the lack of power in the literature (in addition to the small sample sizes) could stem from the significant heterogeneity across studies. This may be an important source of discordant reports, and makes it more difficult to draw conclusions from the literature. The public availability of raw data should greatly enhance our ability to understand neuroanatomical variability and increase our chances of detecting reliable neuroimaging phenotypes for neurodevelopmental disorders such as autism. Towards this aim, all our analysis scripts, our software for quality control, and the list of subjects we included have been made openly available on the Web, to facilitate the replication, critical appraisal and extension of our current results: https://github.com/neuroanatomy/Cerebellum.

In conclusion, we did not find evidence for a difference in cerebellar volume between subjects with ASD and controls, neither in the literature nor in the ABIDE cohort. This result does not rule out a possible involvement of the cerebellum in the aetiology of ASD. We could also imagine that some other measurement of cerebellar anatomy, more sophisticated than simple volume measurements, may be linked to autism in a future. However our current results do not provide evidence to justify a specific focus on the study of the cerebellum instead of any other brain structures. We reached a similar conclusion after a similar analysis of the corpus callosum (Lefebvre et al. 2015), another structure that had traditionally captured the interest of the research community. Based on these experiences, we can only advocate for a broad analysis of all neuroanatomical phenotypes available. For this effort to be successful, our community needs to continue developing the data sharing initiatives that will allow us to increase statistical power, decrease heterogeneity, and avoid the biases preventing researchers from benefiting from the work of each other.

## Acknowledgements

The research leading to these results has received funding from the Fondation de France project “Développement du plissement cortical dans les troubles du spectre autistique: caractérisation morphométrique et génétique”, ERA-Net NEURON Cofund Programme under “Horizon 2020” (SynPathy project), Institut Pasteur, Bettencourt-Schueller Foundation, Centre National de la Recherche Scientifique, University Paris Diderot, Agence Nationale de la Recherche programme “Investissements d’avenir”, Labex BioPsy, ANR10-LABX-BioPsy, ANR-11-IDEX-0004-02, Labex GenMed, ANR SynDiv, Conny-Maeva Charitable Foundation, Cognacq Jay Foundation, Orange Foundation, Fondamental Foundation. The group of LRR is member of the Labex BioPsy and ENP Foundation.

